# Algal Neurotoxin Biosynthesis Repurposes the Terpene Cyclase Structural Fold Into an *N*-prenyltransferase

**DOI:** 10.1101/2020.03.31.014811

**Authors:** Jonathan R. Chekan, Shaun M. K. McKinnie, Joseph P. Noel, Bradley S. Moore

## Abstract

Prenylation is a common biological reaction in all domains of life whereupon prenyl diphosphate donors transfer prenyl groups onto small molecules as well as large proteins. The enzymes that catalyze these biotransformations are structurally distinct from ubiquitous terpene cyclases that instead assemble terpene molecules via intramolecular rearrangements. Herein we report the structure and molecular details of a new family of prenyltransferases from marine algae that repurposes the terpene cyclase structural fold for the *N*-prenylation of glutamic acid during the biosynthesis of the potent neurochemicals domoic acid and kainic acid. We solved the X-ray crystal structure of the prenyltransferase found in domoic acid biosynthesis, DabA, and show distinct active site binding modifications that remodel the canonical Mg^2+^-binding motif. We then applied our structural knowledge of DabA and a homologous enzyme from the kainic acid biosynthetic pathway, KabA, to alter their isoprene donor specificities (geranyl versus dimethylallyl diphosphate) by a single amino acid switch. While the diatom DabA and seaweed KabA enzymes share a common evolutionary lineage, they are distinct from all other terpene cyclases, suggesting a very distant ancestor.

**Significance:** Domoic acid is a neurotoxin produced by marine algae that readily bioaccumulates in shellfish and significantly impacts both human and animal life. The first committed step of the biosynthesis of domoic acid is the *N*-prenylation of L-glutamic acid by the enzyme DabA. By solving the crystal structure of DabA, we demonstrate that this enzyme has repurposed the common terpene cyclase fold to catalyze an extremely unusual reaction, *N*-prenylation of an unactivated primary amine. Application of these structural insights enabled rational engineering of two *N*-prenyltransferase enzymes to accept alternative prenyl donors. Ultimately, these results not only expand the scope of reactions catalyzed by a terpene cyclase family member, but will help inform future domoic acid environmental monitoring efforts.

## Introduction

Marine and freshwater algae produce neurotoxins that encompass a range of structural classes and biological targets (1, 2). During harmful algal blooms (HABs), these neurotoxins are produced and subsequently accumulate in the environment leading to numerous damaging effects including fish and bird mortality, beach and fishery closures, and human toxicity (2, 3). For example, the ionotropic glutamate receptor agonist domoic acid is produced by marine diatoms of the genus *Pseudo-nitzschia* (Scheme 1). Blooms of these diatoms regularly occur around the world, and, as a consequence, human consumption of seafood contaminated with domoic acid leads to Amnesic Shellfish Poisoning, which is characterized by memory loss, seizures, and in severe cases death (4, 5).

Domoic acid is a member of the kainoid family of natural products along with the red macroalgal product kainic acid (Scheme 1). While kainic acid shares a conserved cyclic core with domoic acid, the prenyl moiety is shorter and leads to different bioactive properties. Even though it similarly targets ionotropic glutamate receptors, kainic acid is less potent (6) and has instead been used as both an anthelminthic drug (7, 8) and a chemical reagent to study neurological disorders (9).

Recent work on the biosyntheses of domoic acid and kainic acid established the origin of the prenyl groups in an unusual first committed biosynthetic reaction (10, 11) (Scheme 1). In the case of domoic acid, DabA adds a geranyl group to L-glutamate to generate *N*-geranyl-L-glutamate (NGG), while KabA uses the shorter five carbon dimethylallyl diphosphate prenyl donor during kainic acid biosynthesis to produce the intermediate prekainic acid (Scheme 1). Even though terpene biochemistry is common across all domains of life in the production of bioactive terpenoids, the DabA and KabA *N*-prenylating enzymes are unusual for several reasons. First, the reaction catalyzed by DabA and KabA is uncommon in Nature. Several examples exist of aromatic primary amines, such as those found in adenines, serving as sites of prenylation reactions (12, 13). However, to our knowledge, the only examples of *N*-prenylation of an unactivated primary amine are found in the cyanobactin class of natural products (14, 15). However, cyanobactin prenyltransferases utilize peptidic substrates, while DabA and KabA employ a free amino acid.

In addition to unusual biochemistry, DabA/KabA are also very distinct by amino acid sequence. Querying DabA by BLAST search revealed that the closest homologs outside of those found in kainic biosynthesis have E-values of 10^−4^. A more detailed analysis based on hidden Markov models (HMMs), suggested that both DabA and KabA are distantly related to terpene synthases and more specifically terpene cyclases (16). This initial classification was further supported by structural prediction using both PHYRE2 (17) and I-TASSER (18). The terpene cyclase assignment by both HMM and structural prediction was unexpected as terpene cyclases are not known to catalyze intermolecular prenylation reactions. Instead, the larger terpene synthase family generally catalyzes intramolecular cyclization reactions or loss of the diphosphate moiety of the substrate to yield a linear product (Scheme 1). This type of enzymology is fundamentally different than the intermolecular *N*-prenylation reaction observed in KabA and DabA.

**Scheme 1:**
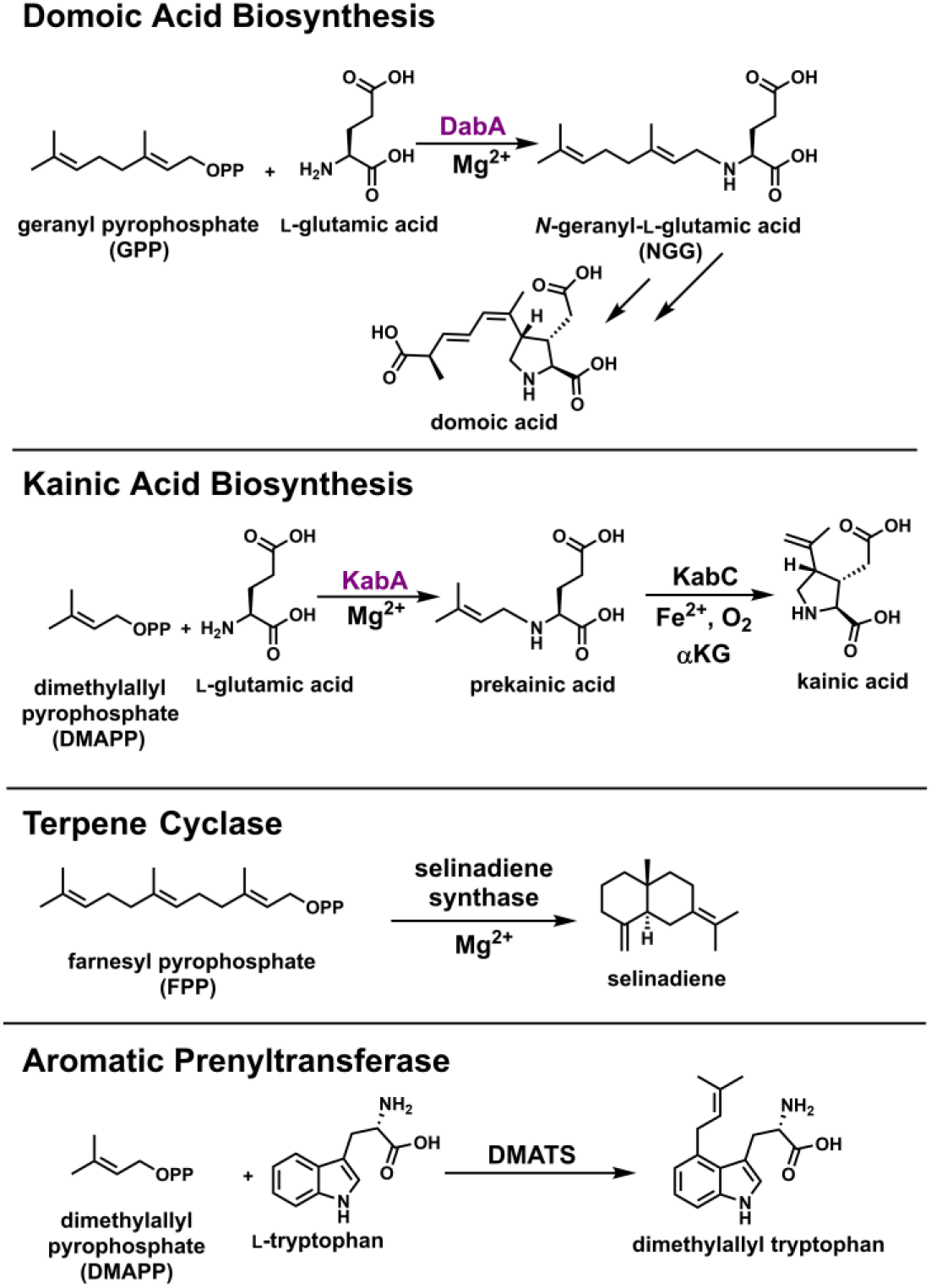
Examples of glutamate *N*-prenyltransferases, terpene cyclases, and prenyltransferases. Dimethylallyl tryptophan synthase is abbreviated as DMATS.

Even though terpene cyclases are not known to catalyze intermolecular prenylation reactions, Nature has evolved a separate class of enzymes called ABBA prenyltransferases that do (Scheme 1) (19). These enzymes utilize prenyl diphosphate substrates and catalyze the transfer of the prenyl group onto a variety of activated functional groups. ABBA prenyltransferases are found in numerous natural product biosynthetic pathways and they are defined by an αββα secondary structural motif (20, 21). This structural arrangement contrasts with the exclusively α-helical fold of terpene cyclases and the one predicted to comprise DabA/KabA.

To better understand the basis for DabA’s unusual activity as a putative member of the terpene cyclase protein family, we obtained a high-resolution crystal structure of DabA in complex with both substrate analogs and products. Validation of our structural observations by mutagenesis helped to reveal how the activate site has been modified to enable prenylation reactions. We further employed this structural information to demonstrate a key residue that assists in dictating prenyl group length specificity and thus helps differentiate the DabA and KabA enzymes from each other.

## Results and Discussion

### Overall Structure

We crystallized and solved the structure of an N-terminally truncated DabA construct in complex with GSPP (geranyl S-thiolodiphosphate) and Mg^2+^ to 2.1 Å resolution. DabA forms a monomer in the asymmetric unit and gel filtration analysis suggests that this is the physiological assembly (Figure S1). Structurally, DabA is composed primarily of an α-helical core that forms a terpene cyclase α-domain, consistent with previous structural prediction (Figure 1A) (10). Analysis by the Dali server (22) indicates strong structural homology to bacterial terpene cyclases and it is most similar to selinadiene synthase from *Streptomyces pristinaespiralis* (2.5 Å r.m.s.d. over 298 aligned Cα) (Figure 1B and Scheme 1) (23). This high level of structural conservation is observed despite low levels of sequence identity (<15% identity) (Figure 1C). While the terpene cyclase core is conserved, DabA has two additional features not observed in typical terpene cyclases. First, DabA contains N- and C-terminal extensions that join together to form a small α-helical bundle (Figure 1B,C). This region connects to the core with a β-hairpin motif. A second insertion is found within the terpene cyclase domain and forms a short helical and β-hairpin extension.

**Figure 1:**
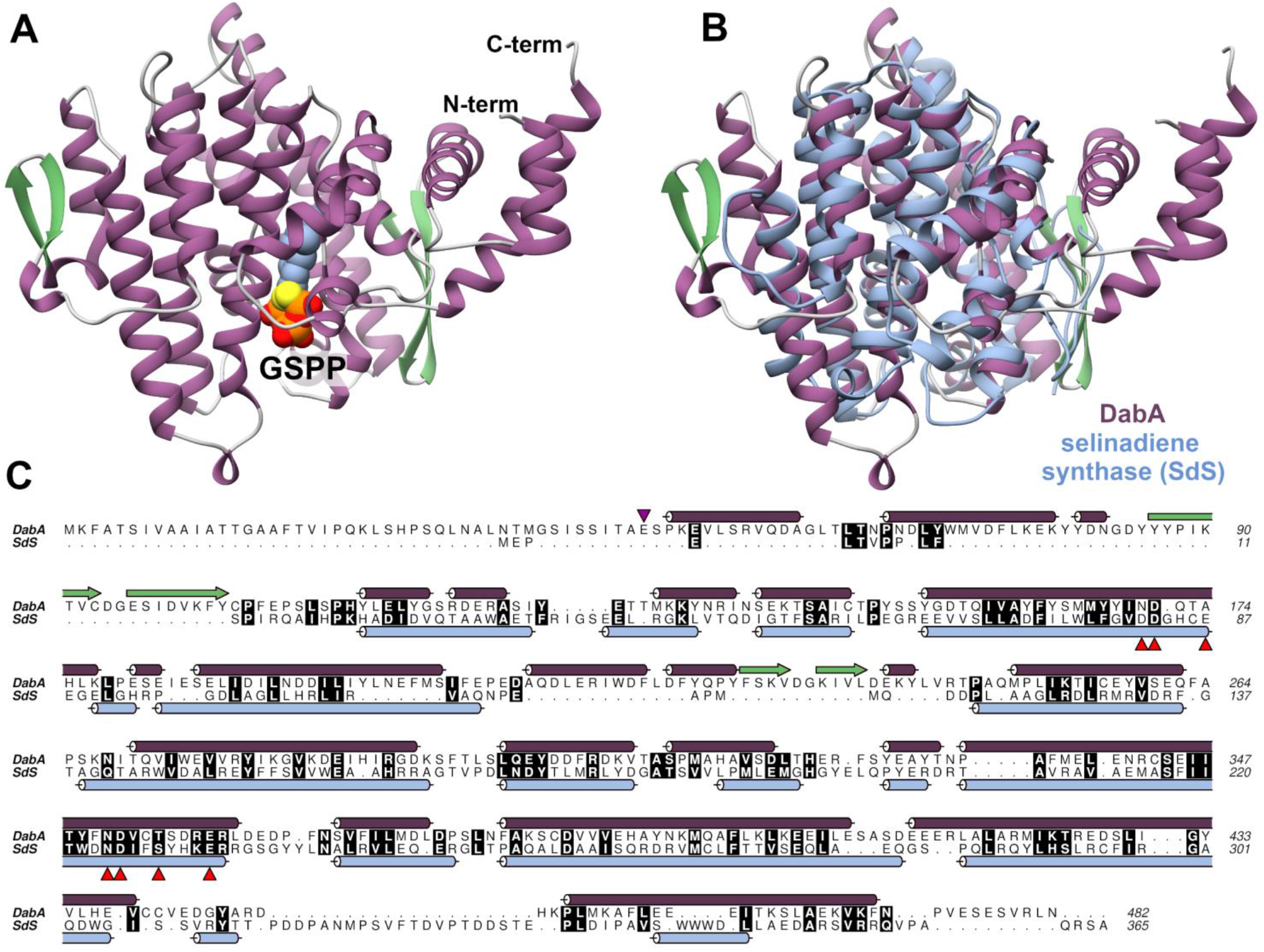
A) Secondary structure of DabA with GSPP also shown. B) Structural alignment of DabA (purple) and selinadiene synthase (blue). C) Sequence alignment of DabA and selinadiene synthase (SdS, PDB accession: 4OKZ). Secondary structure is overlaid next to the sequence and colored according to panel B. The terpene cyclase motifs are indicated by red arrows. The purple arrow indicates the start of the DabA construct used for crystallography.

### Active Site

While the overall structure and location of the active site is conserved between DabA and terpene cyclases, many of the molecular details are very different. Examination of the DabA active site reveals the less-hydrolyzable GPP mimic GSPP bound in at least two distinct conformations (Figure 2A). In both cases, the geranyl group of GSPP stretches into a narrow hydrophobic pocket while the differences between the two conformations lies in the position of the α-phosphate. Two magnesiums are also found in the active site. To give more evidence to the location of these metal ions, DabA crystals were soaked with MnCl_2_ to replace magnesium with the more electron dense and easily identifiable Mn^2+^ divalent cation (Figure S2). The first magnesium coordinates to the α-phosphate in each of the two GSPP conformations along with a water, Asn351, Glu359, and Thr355. The second magnesium is located adjacent to the first and interacts with one of the α-phosphates, three waters, Glu359, and the backbone carbonyl of Phe366. This arrangement contrasts with canonical Class I terpene cyclases which contain two key magnesium binding motifs that typically bind three magnesium ions. The first motif, called the NSE motif, is composed of a NDxxSxxxE sequence. This NSE motif is well conserved in DabA and is composed of Asn351, Thr355, and Glu359 (Figure 1C and 2B). Superimposition of the DabA active site with selinadiene synthase reveals clear alignment of both the residues and magnesium ion (Figure 2C). Notably, DabA also appears to coordinate an additional magnesium ion near the NSE motif not observed in the structure of other terpene cyclases.

**Figure 2:**
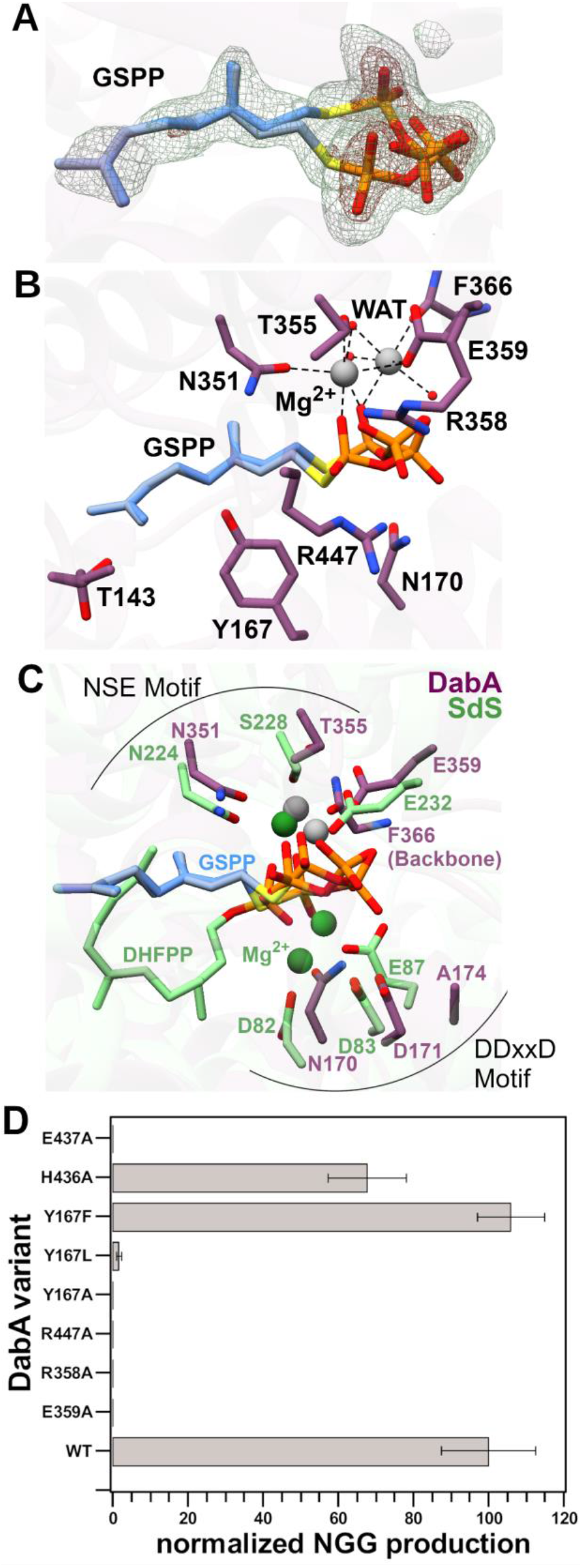
A) Active site of DabA in complex with GSPP, which is shown in two orientations. F_o_-F_c_ maps were generated by removing GSPP and completing refinement without ligand. Meshes are contoured to 3σ (green) and 8σ (red). B) Active site of DabA with important residues highlighted. C) Alignment of DabA (purple) and SdS (green) active sites. DHFPP is 2,3-dihydrofarnesyl diphosphate. D) Activity assays showing the effect of different mutations on the production of NGG by DabA.

The second magnesium binding motif found in terpene cyclases is defined by an aspartate rich DDxxD/E motif. Sequence alignments of DabA with selinadiene synthase show that this motif has been instead mutated to NDxxA and its absence may have contributed to the difficulty in initially identifying DabA as a member of the terpene cyclase family (Figure 1C) (10). Examination of the crystal structure shows that the canonical DDxxD motif has not been replaced by another series of magnesium binding residues; no magnesium is present in this region of the active site (Figure 2C). The active site arrangement of DabA is also distinct from the recently described class IB terpene synthases that are responsible for the biosynthesis of large terpene products (C_25_/C_30_/C_35_) (24). Class IB terpene synthases lack the canonical NSE and DDxxD motifs, but have replaced them with new aspartate rich motifs that occupy the same location within the active site (24). This contrasts with DabA which retains the NSE motif, but lacks the DDxxD motif or any substitute.

Other features prevalent in terpene cyclase substrate binding, such as cationic Arg or Lys residues interacting with the diphosphate moiety, are also well conserved. Specifically, Arg358 and Arg447 are both less than 3.2 Å away from the α-phosphate and likely engage in hydrogen bonding (Figure 2B).

Glutamic acid was present in the crystallization buffer, but it was not observed in the structure. In an effort to determine the location of the glutamate binding pocket, we crystallized DabA in the presence of Mg^2+^ and the NGG reaction product and solved the structure to a resolution of 2.1 Å (Figure 3A). Examination of the active site revealed density for NGG with the geranyl group again extended into the hydrophobic tunnel (Figure S2). Surprisingly, the glutamic acid derived portion of NGG was bound in the same location as the diphosphate end of GSPP. This would suggest a mechanism wherein DabA must first lose diphosphate from the active site before glutamic acid could bind. While possible, we do not favor this mechanism because it requires prolonged stabilization of the carbocation intermediate and protection from hydration in a solvent exposed pocket. Instead, we propose that a small side pocket in the active site could be the binding site for glutamic acid (Figure 3B). In both the NGG and GSPP co-crystal structures, the pocket contains five ordered water molecules and is located near the carbocation intermediate that is predicted to form during the course of the reaction. We evaluated this putative glutamate binding pocket by making mutations to two residues that occupy this putative binding site, Glu437 and His436. Activity assays were completed with both DabA variants (Figure 2D). The Glu437Ala DabA variant was completely inactive, while the His436Ala variant had reduced activity. We further evaluated the His436Ala mutation by obtaining Michaelis-Menten kinetic parameters. Compared to WT DabA, the K_m app_ for glutamic acid increased 3-fold, suggesting that this residue may be involved in glutamate binding (Table 1).

**Figure 3:**
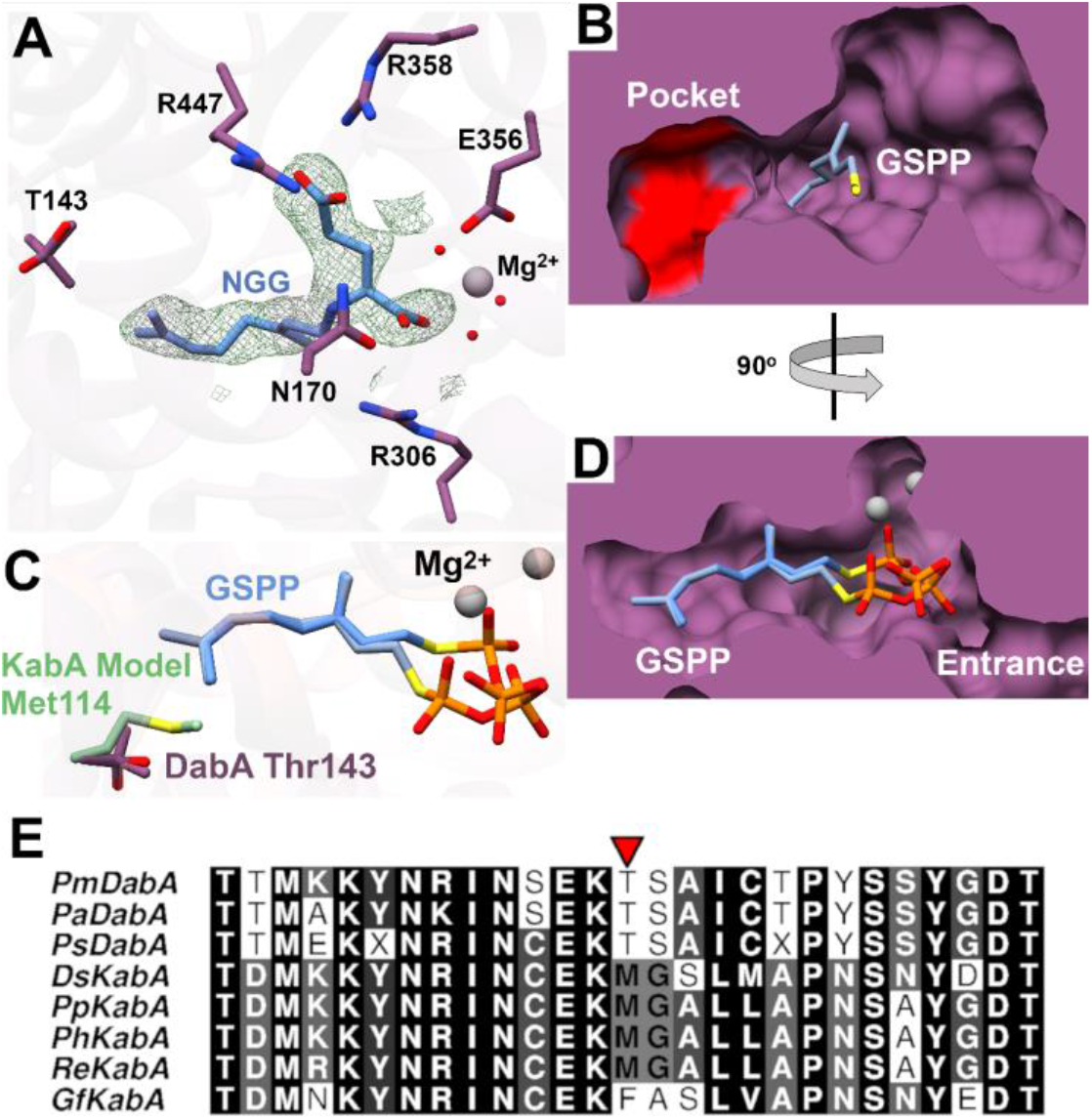
A) Active site of DabA in complex with NGG. F_o_-F_c_ maps were generated by removing NGG and completing refinement without ligand. Mesh was contoured to 3σ (green). B) Surface model of DabA indicates a side pocket is present that may accommodate L-Glu binding. His436 and Glu437 are shown in red. C) Overlay of the DabA active site and a model of KabA. D) Surface modeling of the active site shows a narrow tunnel that accommodates GSPP. E) Sequence alignment of all DabA/KabA homologs. The residue of interest is indicated by a red arrow.

**Table 1:** Michaelis-Menten kinetic constants for DabA, KabA, and amino acid variants.

| Enzyme | GPP |  |  | DMAPP |  |  | L-Glu |  |  |
| --- | --- | --- | --- | --- | --- | --- | --- | --- | --- |
| | $k_{cat\ app}$<br>($\text{min}^{-1}$) | $K_m$<br>($\mu\text{M}$) | $k_{cat\ app}/K_m$<br>($\text{M}^{-1}\text{min}^{-1}$)*$10^6$ | $k_{cat\ app}$<br>($\text{min}^{-1}$) | $K_m$<br>($\mu\text{M}$) | $k_{cat\ app}/K_m$<br>($\text{M}^{-1}\text{min}^{-1}$)*$10^6$ | $k_{cat\ app}$<br>($\text{min}^{-1}$) | $K_m$<br>(mM) | $k_{cat\ app}/K_m$<br>($\text{M}^{-1}\text{min}^{-1}$)*$10^3$ |
| DabA | $4.3 \pm 0.1$ | $0.33 \pm 0.03$ | 12.9 | Non MM | Non MM | | $4.8 \pm 0.3$ | $21 \pm 4$ | 0.23 |
| DabA T143M | $0.65 \pm 0.04$ | $13 \pm 2$ | 0.052 | $1.2 \pm 0.1$ | $130 \pm 20$ | 0.0092 | | | |
| DabA H436A | | | | | | | $6.9 \pm 0.6$ | $68 \pm 16$ | 0.10 |
| KabA | $0.29 \pm 0.01$ | $89 \pm 10$ | 0.0033 | $1.3 \pm 0.1$ | $1.1 \pm 0.2$ | 1.2 | | | |
| KabA M114T | $0.51 \pm 0.02$ | $0.51 \pm 0.06$ | 1 | $0.28 \pm 0.01$ | $2.2 \pm 0.3$ | 0.13 | | | |

### Mutagenesis Supports Active Site Residues

Based on the co-crystal structures, we designed a series of mutations and tested the resulting DabA variants for their ability to catalyze production of NGG (Figure 2D). DabA contains several charged residues in the active site to coordinate both the positively charged Mg^2+^ ions and the negatively charged phosphates. Consistent with this hypothesis, mutation of the Mg^2+^ binding Glu359 residue to Ala completely abolishes activity. Similarly, mutation of either Arg358 or Arg447 to Ala results in an inactive DabA variant. We next explored the significance of Tyr167 by creating a series of mutations. This residue is located directly below GSPP in the co-crystal structure and could be responsible for different roles in catalysis including stabilizing the carbocation, maintaining the shape of the hydrophobic tunnel, or deprotonating the α-amine of the glutamic acid co-substrate. Mutation of Tyr167 to Phe resulted in no change in activity, indicating that Tyr167 does not serve as a catalytic base. Instead, the DabA Tyr167Leu and Tyr167Ala variants exhibited dramatically reduced or no activity, respectively. This indicates that Tyr167 is important in shaping the tunnel as the much smaller Tyr167Ala residue appeared completely inactive under the reaction conditions, while the slightly smaller Tyr167Leu retained some activity. Furthermore, the observation that only phenylalanine can successfully substitute for tyrosine is consistent with a carbocation stabilizing activity for Tyr167. Terpene cyclases often employ aromatic residues in the active site and both crystal structures and modeling studies have suggested that they are responsible for stabilizing the carbocation of the mechanistic intermediate through the use of cation-π interactions (25–27).

### DabA Kinetic Analysis

Based on the unusual activity of DabA as a structural member of the terpene cyclase family, we sought to determine the kinetic parameters of DabA and compare them to both canonical terpene cyclases and prenyltransferases (Figure S3). WT DabA used the native GPP substrate with a K_m app_ of 0.33 μM and k_cat app_ of 4.3 min^−1^. These constants are similar to those found in other characterized prenylating or cyclizing enzymes. Specifically, the low K_m app_ for the prenyl donor is similar to the sub-micromolar values observed in structurally similar terpene cyclase class of enzymes, such as geosmin synthase (28), pentalenene synthase (29), and aristolochene synthase (30). Moreover, DabA has turnover numbers similar to slower members of the terpene cyclase family (28, 30) as well as ABBA aromatic prenyltransferases such as FtmPT2 (31).

In addition to GPP, kinetics parameters were also determined for L-glutamic acid and a k_cat app_ of 4.8 min^−1^ was observed, consistent with the value obtained for GPP. The K_m app_ for L-glutamic acid was 21 mM, much higher than the K_m app_ of GPP. Enzymes are typically observed to possess K_m app_ values within 10-fold of the physiological concentration of substrates (32, 33). DabA is expected to be localized to the diatom plastid (10), and glutamic acid is amongst the most concentrated metabolites in that organelle, with values of 14 mM (34) and 74 mM (35) reported from different plants. Therefore, even though the K_m app_ value for glutamic acid is high, it is well aligned with in vivo concentrations.

### Basis for prenyl group selectivity

The major structural difference between the two marine kainoid natural products, domoic acid and kainic acid (Scheme 1), is the biosynthetic origin of the prenyl donor, either GPP or DMAPP, respectively (10, 11). The co-crystal structure of DabA with GSPP indicated that the carbon chain is positioned within a hydrophobic tunnel (Figure 3D). We hypothesized that alterations to the length of the tunnel would be a major factor in the selectivity of DabA for GPP and KabA for DMAPP. Therefore, we created a model of the *Digenea simplex* KabA structure using the I-TASSER server (18) and DabA as a template. Overlay of the DabA structure and KabA model shows the active site is largely conserved (Figure S4). Examination of the hydrophobic tunnel revealed a threonine to methionine substitution in KabA (Figure 3C). Sequence alignments of the known DabA and KabA enzymes suggest that the identity of this residue can predict the prenylation activity (Figure 3E). The DabA homologs found in diatoms all contain a threonine, while the red algal kainic acid biosynthetic KabAs have a larger residue, either methionine or phenylalanine.

We evaluated the significance of DabA Thr143 by completing Michaelis-Menten kinetics of DabA, KabA, and variants wherein this tunnel residue was switched (Figure S3). Exchange of the native threonine or methionine tunnel residue in both DabA and KabA dramatically decreased catalytic efficiency for the native substrate and instead promoted the use of the non-native substrate (Table 1). Specifically, the DabA Thr143Met mutation decreases catalytic efficiency for GPP by ~250-fold. WT DabA did not display canonical Michaelis-Menten kinetics with DMAPP, but had a maximum rate of 0.6 min^−1^ at a DMAPP concentration of 500 μM. Conversely DabA Thr143Met variant does display typical kinetics with a k_cat app_ of 1.2 min^−1^ and K_m app_ of 130 μM.

A similar analysis was completed for KabA to evaluate the effect of Met114 on substrate specificity. Similar to DabA, the KabA Met114Thr mutation decreases catalytic efficiency for DMAPP by 10-fold and increases catalytic efficiency for GPP by ~300-fold. In both cases, not only is the native substrate selected against, but activity with the non-native substrate improves.

This “molecular ruler” basis for substrate selectivity has been observed in other prenylating enzymes. For example, FPP synthase is an iterative enzyme that first produces GPP from DMAPP and IPP. It then uses GPP and IPP to generate FPP. Constriction of the hydrophobic pocket with bulky residues biases formation of GPP over FPP (36, 37), while expansion of the pocket promotes prenylation of FPP to generate C_20_ and C_25_ products (38). In another case, the tyrosine prenyltransferase PagF natively utilizes DMAPP but a single point mutation can expand the hydrophobic binding site to generate a PagF variant that instead favors GPP as a substrate (39).

### Alternative Substrate Selectivity

The kinetic analysis and mutagenesis revealed that both DabA and KabA are selective for their respective prenyl donor and exhibit greatly reduced catalytic efficiencies when the non-native prenyl donor is used. To further explore the substrate scope, we tested both DabA’s and KabA’s ability to utilize alternative substrates. We began by synthesizing a suite of prenyl diphosphate derivatives: propargyl, 2-butynyl, allyl, crotyl, and farnesyl. Each of these was evaluated as a potential substrate for DabA and KabA and monitored by LC-MS (Figure S5). Both DabA and KabA exhibited high levels of selectivity with neither enzyme utilizing the alkyne-containing prenyl donors nor the allyl diphosphate substate. From a mechanistic standpoint, these substrates would not be able to readily stabilize the carbocation that forms upon loss of the diphosphate because no tertiary carbocation could be formed. The crotyl diphosphate, however, would form a secondary carbocation, which is more favorable than a primary carbocation. Only KabA, and not DabA, could use this substrate to form the crotyl-L-glutamate product. Finally, we tested the 15-carbon farnesyl diphosphate substate and found that DabA alone could form the farnesyl-L-glutamate product.

DabA was also tested for its ability to utilize alternative divalent cations as cofactors (Figure S6). DabA activity was highest when magnesium was used as the cofactor and no product was observed when a divalent cation was omitted from the reaction conditions. Manganese could substitute for magnesium, but led to ~28% lower activity. Other metals were poor replacements for magnesium with nickel and zinc reducing DabA activity by 69% and 96%, respectively, as compared to magnesium. Finally, calcium appeared unable to successfully serve as a cofactor.

### Gene Phylogeny

While the structure of DabA shows high similarity to bacterial terpene cyclases, BLASTp analysis of sequences in the NCBI protein database reveals only distant homologs outside of kainic acid biosynthetic genes, with the closest homolog having a blast E-value of higher than 10^−4^. We therefore aimed to determine the evolutionary history of these glutamate *N*-prenyltransferases by constructing a maximum likelihood phylogenetic tree. Due to the low sequence similarity of DabA/KabA to any publicly available protein sequence, we decided to generate a global view of the entire terpene cyclase family (InterPro families IPR034741 and IPR034686). Additionally, we seeded in the characterized red algae terpene cyclases (40, 41) along with the closest DabA sequence homologs identified from PSI- and DELTA-BLAST (Table S3).

The tree forms two distinct halves with one composed of plant proteins and the other a mixture of bacterial, algal, and fungal proteins (Figure 4). This tree correlates with the α/β-domain organization of many plant terpene cyclases and the lone α-domain of the enzymes from the remaining organisms. The DabA proteins from *Pseudo-nitzschia* spp. diatoms cluster together and are closely related to the red algal KabA proteins. This *N*-prenyltransferase protein region is most related to the bacterial, red algal, and fungal enzymes. Similarly, the four characterized red algal terpene cyclases form a distinct region within the bacterial branch, consistent with previous phylogenetic analysis (40). Even though both KabA and the red algae terpene cyclase sequences are found associated with the microbial clade, the distance between these proteins suggests that the kainoid biosynthetic genes did not simply arise from algal terpene cyclases through a recent gene duplication event. Instead, a more distant evolutionary relationship is likely. Furthermore, the phylogeny of the diatom DabA and the red algal KabA proteins does not match the evolution of the organisms themselves. Diatoms are not closely related to red algae and are thought to have originated from a secondary endosymbiotic event from an ancestral red alga (42). Instead, red algae are more closely related to green plants (43). This inconsistency again suggests a curious evolutionary history that may employ mechanisms such as horizontal gene transfer.

**Figure 4:**
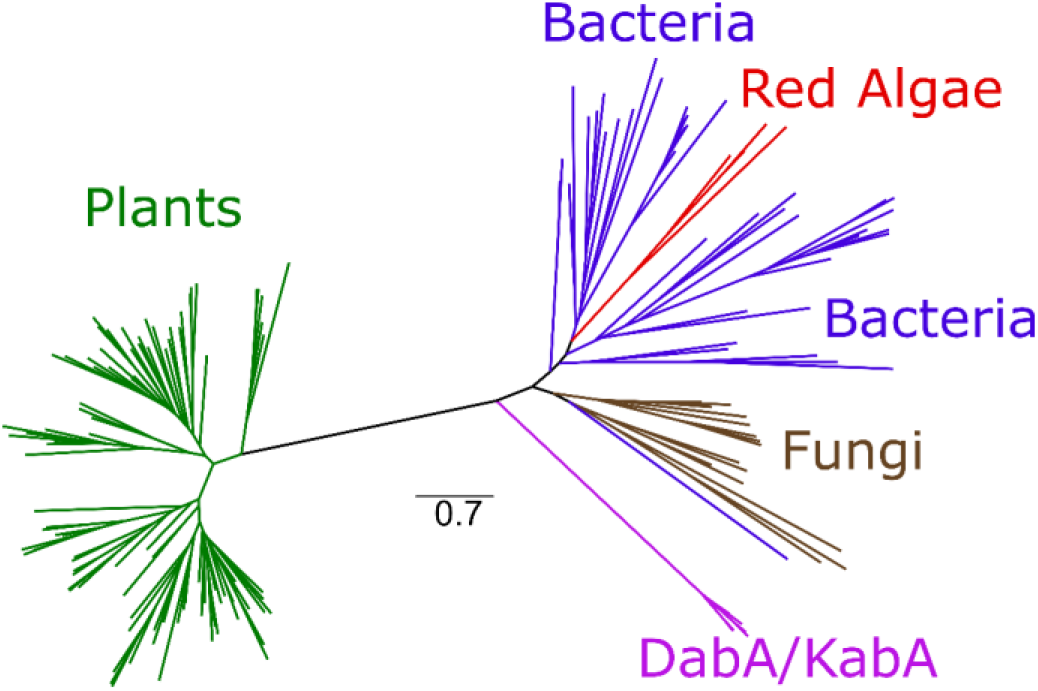
A representative selection of all terpene cyclases family members was used to generate a maximum likelihood phylogenetic tree.

## Conclusion

The biosynthesis of the toxic algal natural products domoic acid and kainic acid begins with an uncommon *N*-prenylation reaction of a free amine. Our work demonstrated that DabA and KabA accomplish this reaction by repurposing the prevalent terpene cyclase fold. Instead of an intramolecular cyclization reaction, DabA and KabA perform an intermolecular prenylation reaction, an activity not previously observed in this family. Our structural work revealed that DabA uses the same active site as a canonical terpene cyclase, but has employed two key modifications. First, the active site has been narrowed to create a hydrophobic tunnel that prevents cyclization. Second, we have proposed a small side pocket that can accommodate the glutamate co-substrate.

Our results also help explain the structural differences between DabA and KabA that dictate substrate specificity. By creating a single point mutation that changes the length of the tunnel, we were able to successfully alter the specificity of this family of *N*-prenyltransferases. Moreover, this residue can serve as an indicator for the physiological prenyl substrate of an unknown homolog. Finally, phylogenetic analysis reinforces the unusual evolutionary history of these enzymes first suggested by simple amino acid sequence alignments. Instead of being related to plant or red algal enzymes, they form their own distinct branch. This affiliation is in spite of the distant relationship between red algae and diatoms. Examination of hypothetical homologs from the acromelic acid producing mushroom may enable better insight into how these enzymes evolved (44). In addition, the detailed understanding of DabA described here may help inform the creation of PCR based assays that can detect DabA expression in the environment and serve as an early warning system before domoic acid accumulates to toxic levels. This approach is analogous to the successful methods used in freshwater neurotoxin monitoring (45). Finally, our results reinforce the observation that understudied organisms create opportunities to discover new types of enzyme biocatalysts (46).

## Materials and Methods

SI Appendix includes methods for expression, purification, and mutagenesis of DabA and KabA. Methodologies for crystallization, activity assays, kinetics, and phylogenetics are also detailed. Finally, synthetic procedures and small molecule characterization are available in the SI Appendix.

### Data deposition

The structure factors and coordinates have been deposited in the Protein Data Bank, www.wwpdb.org: DabA in complex with Mg^2+^ and GSPP (PDB ID code 6VKZ), DabA in complex with Mn^2+^ and GSPP (PDB ID code 6VL0), and DabA in complex with Mg^2+^ and NGG (PDB ID code 6VL1). The NCBI accession numbers for proteins discussed in this work are: AYD91073.1 (DabA) and QCC62379.1 (KabA).

## Author Contributions

J.R.C., S.M.K.M, J.P.N., and B.S.M. designed the study. J.R.C. performed experiments. S.M.K.M synthesized compounds. J.R.C. and B.S.M. wrote the manuscript with input from all authors.

## Notes

The authors declare no competing financial interests.

## Acknowledgement

We thank G. Louie (Salk Institute) and the staff of the ALS at beamlines 8.2.1 and 8.2.2 (Berkeley, California) for assistance with data collection, G. Rouse (Scripps Institution of Oceanography) for helpful discussions, and D. Nguyen for assistance with synthesis.

