## Supplemental Information for "Algal Neurotoxin Biosynthesis Repurposes the Terpene Cyclase Structural Fold Into an *N*-prenyltransferase"

**Table of Contents**

1. **Biological Methods ........................................................................ S2**

2. **Synthetic Methods ........................................................................ S5**

3. **Supplementary Tables**
   Table S1. Primers used in this study ........................................ S7
   Table S2. Crystallographic refinement statistics ........................ S8
   Table S3. Sequences used in phylogenetic tree ........................ S9

4. **Supplementary Figures**
   Figure S1. Gel filtration analysis of DabA ........................................ S10
   Figure S2. DabA crystal structure with $Mn^{2+}$ ........................................ S11
   Figure S3. Michaelis-Menten kinetic analysis of DabA and KabA ............ S12
   Figure S4. Stereoview of the DabA and KabA active site alignment ........ S13
   Figure S5. LC-MS analysis of alternative prenyl donors ........................ S14
   Figure S6. DabA activity assays with alternative cations ........................ S17

5. **Supplementary References** ........................................................ S18

### Biological Methods

##### Cloning and Mutagenesis

PCR of genes was completed using PrimeStar HS DNA polymerase (Takara Bio), while the vector backbone was amplified using PrimeStar Max (Takara Bio). Protein expression constructions were generated in the pET28a vector using the NEBuilder HiFi DNA Assembly mix. Mutations were generated by a three-piece assembly using the vector, an N-terminal fragment containing the mutation, and a C-terminal fragment containing the mutation. Plasmids were purified from *Escherichia coli* DH5$\alpha$ cells and inserts were Sanger sequenced to confirm their identity. Vectors were transformed into *E. coli* BL-21 cells for subsequent expression. The DabA protein was constructed as Ser26 to C-term truncation to remove the N-terminal signal peptide. All subsequent DabA variants were based on this construct.

##### Protein Expression and Purification

*E. coli* BL-21 cells harboring the appropriate vector were grown overnight in 10 mL of lysogeny broth supplemented with 50 $\mu$g/mL kanamycin. A portion of the overnight culture (4 mL) was used to inoculate 1 L of terrific broth supplemented with 50 mg/L kanamycin in a 2.4 L Erlenmeyer flask. The flasks were placed in a 37 $^{o}$C shaking incubator until the $OD_{600}$ reached approximately 1. The incubator was then cooled to 18 $^{o}$C, and the flasks were incubated for an additional 1 h. IPTG (0.5 mM final concentration) was added to induce protein expression, and the flasks were incubated for 18 h. Cells were harvested by centrifugated at 8,000 x g for 10 minutes and resuspended in buffer containing 500 mM NaCl, 20 mM Tris pH 8.0, and 10% glycerol. Cell pellets were frozen and stored at -70 $^{o}$C.

Cells were lysed by sonication with a Qsonica 6 mm tip at 40% amplitude for 12 cycles of 15 seconds on and 45 seconds off. The lysate was centrifuged at 14,000 x g for 30 minutes to remove any insoluble debris. The supernatant was loaded onto a 5 mL HisTrap column (GE Healthcare) and washed with buffer A (1 M NaCl, 20 mM Tris pH 8.0, 30 mM imidazole). Protein was eluted using a linear gradient from 0-100% buffer B (1 M NaCl, 20 mM Tris pH 8.0, 250 mM imidazole) over 40 mL while collecting 5 mL fractions. Fractions were evaluated for purity by SDS-PAGE, and pure fractions were collected.

In the case of DabA used for crystallography, 100 U of thrombin were added to the pooled fractions immediately after purification by the HisTrap column, and the mixture was allowed to incubate for 18 h at 4 $^{o}$C. Complete cleavage of the N-terminal $His_6$ tag was confirmed by SDS-PAGE. Protein was subsequently purified by size exclusion chromatography using a HiLoad 16/60 Superdex 75 prep grade column (GE Healthcare Life Sciences) pre-equilibrated with 100 mM KCl and 20 mM HEPES pH 7.5. Protein containing fractions were collected and concentrated with an Amicon Ultra-15 30kDa centrifugal filter (Millipore Sigma) to approximately 15 mg/mL before aliquoting, freezing on dry ice, and storage at -70 $^{o}$C.

If protein was not to be used for crystallography, pure fractions were directly taken after the HisTrap column and concentrated with an Amicon Ultra-15 30 kDa centrifugal filter (Millipore Sigma). The protein was further purified by size exclusion chromatography using a HiLoad 16/60 Superdex 75 prep grade column (GE Healthcare Life Sciences) pre-equilibrated with 300 mM NaCl, 10% glycerol, and 20 mM Tris pH 8.0. Protein containing fractions were collected and concentrated to approximately 15 mg/mL before aliquoting, freezing on dry ice, and storage at -70 $^{o}$C.

##### Crystallography, Data Collection, Structure Determination, and Structure Refinement

DabA was screened using commercially available sparse matrix screens. Initial crystals were observed in the Wizard III screen (Rigaku Reagents), but could not be successfully optimized. Based on secondary structure prediction and conservation, an N-terminal truncation was made creating a DabA Glu46 to C-term construct. This truncated protein was successfully optimized with hanging drop vapor diffusion as follows: 4 mg/mL protein was preincubated with 2 mM GSPP, 5 mM $MgCl_2$, and 50 mM L-Glu and mixed 1 μL : 1 μL with a mother liquor containing 1.5 M sodium citrate and 0.05 M HEPES pH 7.5. Crystals were grown over a well of 150 μL of mother liquor and began to appear after one month at 9 $^oC$. Crystals were transferred to fresh mother liquor supplemented with 2 mM GSPP, 5 mM $MgCl_2$, and 100 mM L-Glu for 24 h. Immediately prior to vitrification in $LN_2$, crystals were briefly soaked in mother liquor supplemented with 30% D-glucose, 2 mM GSPP, 5 mM $MgCl_2$, and 100 mM L-Glu. Manganese containing crystallized were generated in a similar manner except $MgCl_2$ was replaced with $MnCl_2$ and a 3 h soak was used instead of a 24 h soak. For the NGG containing crystals, the mother liquor was composed of 1.35 M sodium citrate tribasic, 0.05 M HEPES pH 7.5, 5mM $MgCl_2$ and 2 mM NGG and DabA was used at a concentration of 8 mg/mL. NGG containing crystals were not soaked in fresh mother liquor and instead were briefly soaked in mother liquor supplemented with 30% D-glucose, 2 mM NGG, and 5 mM $MgCl_2$ prior to vitrification. All data were collected at the Advanced Light Source Macromolecular Crystallography beamline (8.2.1).

Diffraction data were indexed and scaled using autoPROC (1). Initial phases were determined by using a combination of SAD and molecular replacement. Specifically, crystals were soaked for 3 h in 1 mM methyl mercury chloride prior to vitrification. An initial DabA structure was generated from the weak anomalous signal found in the data set along with PDB:5NX5 using MR-SAD in the Phenix software package (2). This low quality initial structure was used as the search model for PHASER (3) to obtain improved phases. A second model was built using a combination of Buccaneer (4) and Phenix autoBuild (2). The structure was improved by iterative manual adjustments in COOT (5) and refinement with REFMAC5 (6). All collection and refinement statistics are found in Table S2.

##### $d_5$-NGG and $d_5$-PKA Production

$d_5$-NGG and $d_5$-PKA were produced as internal standards to accurately normalize NGG and PKA enzymatic production between LC-MS runs. Both deuterium labeled compounds were produced enzymatically using either native DabA or KabA. A 300 μL solution containing 100 mM L-glutamic acid (2,3,3,4,4-D5, 97-98%, Cambridge Isolate Laboratories), 1 mM DMAPP/GPP, 5 mM $MgCl_2$, 300 mM NaCl, 20 mM Tris pH 8.0, and 25 μM enzyme was incubated at 21 $^oC$ for 18 h. The reaction was filtered, diluted, aliquoted, and stored at -70 $^oC$ for later use.

##### Activity Assays

Activity assays were performed to test the effects of different mutations. Assays were completed in 50 μL total volume and contained 5 nM DabA variant, 5 mM $MgCl_2$, 25 mM L-glutamate pH 8.0, 100 μM GPP, 300 mM NaCl, and 20 mM Tris pH 8.0. Reactions were initiated with enzyme and, after 10 minutes, were quenched with 50 μL of 2% aqueous formic acid containing deuterium labeled NGG. Samples were analyzed using a Bruker amaZon Ion Trap and Agilent 1200 LC-MS with a Synergi Polar-RP 4μ 250 x 4.6 mm column at 0.75 mL/min column using the following method: 0% B (4.5 min), 0 to 5% B (0.5 min), 5 to 26% B (9 min), 26 to 80% B (9 min), 80 to 100% B (1 min), 100% B (1.5 min), 100 to 0% B (2.5 min), 0% B (2 min), wherein A = 0.1% aqueous formic acid, and B = 0.1% formic acid in acetonitrile. Extracted ion chromatograms for the expected NGG product mass, $[M-H]^{-1}$ = 282.2, were integrated. The peak areas were all normalized using the deuterium labeled NGG present in the quenching solution. Each area was then normalized to the amount produced by DabA WT. Assays for each DabA variant were completed in triplicate.

Activity assays were completed to test the ability of DabA and KabA to utilize different prenyl donors. Assays were completed in 100 μL total volume and contained 2 μM DabA variant, 5 mM $MgCl_2$, 100 mM L-glutamate pH 8.0, 200 μM prenyl donor, 300 mM NaCl, and 20 mM Tris pH 8.0. Reactions were initiated with enzyme and, after 4.5 hour, were quenched with 100 μL of 2% aqueous formic acid. The majority of samples were analyzed for product formation by LC-MS using the method described for the mutagenesis assays. For the longer geranyl and farnesyl diphosphate substrate assays, the following method was used: 10% B (0.5 min), 10 to 100% B (19 min), 100% B (1.5 min), 100 to 10% B (2.5 min), 10% B (2 min), wherein A = 0.1% aqueous formic acid, and B = 0.1% formic acid in acetonitrile. The expected product mass was extracted for each assay.

Activity assays were completed to compare the ability of DabA to utilize different divalent cations as cofactors. Assays were completed in 50 μL total volume and contained 5 nM DabA, 5 mM of the chloride salt of the tested cation, 100 mM L-glutamate pH 8.0, 100 μM GPP, 300 mM NaCl, and 20 mM Tris pH 8.0. Reactions were initiated with enzyme and, after 10 minutes, were quenched with 50 μL of 2% aqueous formic acid containing deuterium labeled NGG. Samples were analyzed for NGG production using the LC-MS procedure discussed in the kinetic assay protocol. Each run was integrated for NGG and $d_5$-NGG as described in the kinetic assay protocol and subsequently normalized to the $d_5$-NGG intensity. Assays were completed in quadruplicate and normalized to DabA catalyzed NGG production in the presence of $MgCl_2$.

**Kinetic Assays**

Individual assay conditions were optimized for each enzyme and substrate tested. In general, assays were completed on the 200 μL scale using 5-25 nM enzyme, 5 mM $MgCl_2$, 300 mM NaCl, and 20 mM Tris pH 8.0. For assays measuring kinetics of DMAPP or GPP, L-glutamate pH 8.0 was used at a constant concentration of 100 mM. For assays measuring kinetics of L-glutamate, DMAPP or GPP was used at a concentration of 100 μM. Reactions were initiated with the addition of enzyme and 50 μL was removed at 5, 10, and 15 minutes and quenched in 50 μL of 2% aqueous formic acid containing a standard amount of either $d_5$-NGG or $d_5$-PKA. Each reaction was analyzed using Agilent 1260 infinity HPLC and Bruker amaZon ion trap mass spectrometer LC-MS with a Phenomenex Luna 5μm $C_{18(2)}$ 100 Å 100 x 4.6 mm column using a 40 μL injection. For LC-MS analysis, Solvent A was water + 0.1% formic acid and Solvent B was acetonitrile + 0.1% formic acid.

The method to quantify NGG was: 20% Solvent B for 1.5 minutes; 20 to 100% Solvent B over 4.5 minutes; 100% Solvent B for 1 min; 100 to 20% Solvent B over 1 min; and 20% Solvent B for 3 min. Targeted MS/MS on $[M-H]^-$ of 282.2 *m/z* and 287.2 *m/z* with a width of 4 *m/z* was used to select for NGG and d5-NGG respectively. The chromatogram for each run was extracted for the major $MS^2$ fragment (264.0 *m/z* and 267.8-269.2 *m/z* for NGG and $d_5$-NGG, respectively) and integrated. Peak areas were normalized to the deuterium labeled internal standard. A standard curve of NGG containing the $d_5$-NGG internal standard was generated and used to quantify NGG production over the course of the reaction.

The method to quantify PKA was 2% Solvent B for 1.5 minutes; 2 to 100% Solvent B over 4.5 minutes; 0% Solvent B for 1 min; 0 to 98% Solvent B over 1 min; and 2% Solvent B for 3 min. Targeted MS/MS on $[M-H]^-$ of 216.1 *m/z* and 221.1 *m/z* with a width of 4 *m/z* was used to select for PKA and $d_5$-PKA, respectively. The chromatogram for each run was extracted for the major $MS^2$ fragment (147.6 *m/z* and 152.6 *m/z* for PKA and $d_5$-PKA, respectively) and integrated. Peak areas were normalized to the deuterium labeled internal standard. A standard curve of PKA containing the $d_5$-PKA internal standard was generated and used to quantify NGG production over the course of the reaction.

All data points were completed in triplicate and data fit to the Michaelis-Menten equation using linear regression.

**Phylogenetic Tree**

Efforts to construct a focused phylogenetic tree to evaluate the evolutionary history of KabA and DabA were frustrated by the lack homologs on the amino acid level. Therefore, to construct a more global view of the placement of DabA/KabA within the terpene cyclase family, the entire InterPro families (7) for terpene cyclase-like 2 (IPR034686) and terpene cyclase-like 1, C-terminal domain (IPR034741) were chosen comprising a total of 9,125 sequences. To select representative members for tree construction, the sequences were cluster by amino acid sequence similarity with CD-HIT (8) using a cut off of 50% identity. A member from the 150 largest clusters was selected for inclusion in the tree. In addition to these sequences, the closest DabA/KabA homologs as determined by BLAST (TPR00180.1 and SCL16690.1) along with characterized red algal terpene cyclases (AXN72983.1, AZO92733.1, AZO92734.1, and ASV63464.1) were also added to the sequence list. Amino acid sequence alignment was completed with MAFFT (9) using the default parameters. A maximum likelihood tree was constructed with IQ-TREE(10) using the LG+R6 substitution model. This tree showed DabA/KabA separated from red algal terpene cyclases. To rigorously evaluate this observation, a constrained tree was created that forced the DabA/KabA proteins into the red algal branch. This tree was then evaluated using the approximately unbiased test (11) as implemented in IQ-TREE. A p-value of 0.015 indicated that the constrained tree can be rejected and the DabA/KabA branch is phylogenetically distinct from red algal terpene cyclases.

### Synthetic Methods

GPP (12), DMAPP (13), GSPP (14), and NGG (15) were chemically synthesized using established protocols and matched previous literature characterization.

Alternative organic pyrophosphate molecules were synthesized by adapting literature protocols (12). To a solution of tris (tetrabutylammonium) pyrophosphate (12) (0.5 g, 0.55 mmol) in dry acetonitrile (5 mL) at -35 °C was added alkenyl/alkynyl halide over 2 minutes. The reaction mixture was stirred at -35 °C for 10 minutes, then warmed to room temperature and stirred for 2 hours. The solvent was removed *in vacuo* and resuspended in minimal ion exchange buffer (25 mM $NH_4HCO_3$ in 2% aqueous isopropanol), then passed through 10 mL of DOWEX AG50W-X8 resin ($NH_4$ form), collecting the first 2 column volumes. The material was lyophilized, generating an off white solid, which was used without further purification.

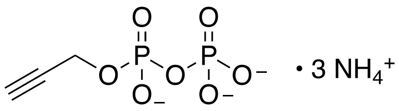

propargyl pyrophosphate trisammonium salt: $^1$H-NMR (500 MHz, $D_2O$ + 0.1% $CH_3OH$) $\delta$ 4.58 – 4.55 (m, 2H, -OC$\underline{H_2}$), 2.88 (t, *J* = 2.4 Hz, 1H, CC-$\underline{H}$); $^{13}$C-NMR (125 MHz, $D_2O$ + 0.1% $CH_3OH$) $\delta$ 80.3, 76.1, 54.2 (d, *J* = 4.0 Hz).

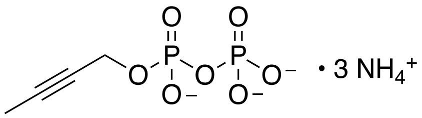

but-2-ynyl pyrophosphate trisammonium salt: $^1$H-NMR (500 MHz, $D_2O$ + 0.1% $CH_3OH$) $\delta$ 4.55 – 4.50 (m, 2H, -OC$\underline{H_2}$), 1.85 – 1.84 (m, 3H, C≡C-$\underline{CH_3}$); $^{13}$C-NMR (125 MHz, $D_2O$ + 0.1% $CH_3OH$) $\delta$ 85.0, 75.5, 55.3, 3.3.

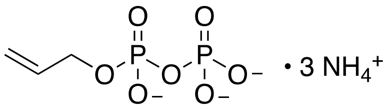

allyl pyrophosphate trisammonium salt: $^1$H-NMR (500 MHz, $D_2O$ + 0.1% $CH_3OH$) δ 6.00 (ddt, 1H, *J* = 17.9, 10.6, 7.4 Hz, $-C\underline{H}=CH_2$), 5.38 (dd, 1H, *J* = 17.3, 1.8 Hz, $-CH=C\underline{H_2}$), 5.25 – 5.19 (m, 1H, $-CH=C\underline{H_2}$), 4.44 (dt, *J* = 7.5, 2.7 Hz, $-OC\underline{H_2}$); $^{13}$C-NMR (125 MHz, $D_2O$ + 0.1% $CH_3OH$) δ 135.0, 117.5, 67.3 (d, *J* = 5.2 Hz).

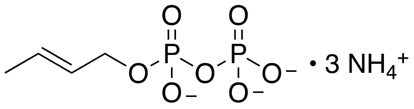

crotyl pyrophosphate trisammonium salt: $^1$H-NMR (500 MHz, $D_2O$ + 0.1% $CH_3OH$) δ 5.91 – 5.80 (m, 1H, $C\underline{H}=CHCH_3$), 5.74 – 5.60 (m, 1H, $-CH=C\underline{H}CH_3$), 4.43 – 4.33 (m, 2H, $-OC\underline{H_2}$), 1.70 (d, *J* = 6.6 Hz, $=CH\underline{CH_3}$); $^{13}$C-NMR (125 MHz, $D_2O$ + 0.1% $CH_3OH$) δ 131.7, 127.1, 67.5, 17.7.

| Primer Name | Sequence |
|---|---|
| DabA S26 p28 NdeI F | GGTGCCGCGCGGCAGCCATATGATGTCGCACCCAAGCCAGCTCAATGCC |
| DabA E46 p28 NdeI F | GGTGCCGCGCGGCAGCCATATGGAAAGTCCGAAGGAAGTTCTCTCCCGC |
| DabA p28 HindIII Rev | GCTCGAGTGCGGCCGCAAGCTTTCAATTGAGGCGAACGGACTCAGACTCAACCGG |
| pET28 Up | CATATGGCTGCCGCGCGGCACC |
| pET28 Down | CT CGAGCACCACCACCACCACCACTGAG |
| DabA T119M For | ACAACCGGATTAACAGCGAGAAAATGTCAGCGATTTGCACC |
| DabA T119M Rev | GGTGCAAATCGCTGACATTTTCTCGCTGTTAATCCGGTTGT |
| DabA E335A For | GTGCACTAGCGATAGGGCAAGATTGGACGAAGATC |
| DabA E335A Rev | GATCTTCGTCCAATCTTGCCCTATCGCTAGTGCAC |
| DabA Y143F For | CGCATATTTTTACTCGATGATGTTCTATATCAATGATCAAACTGCTC |
| DabA Y143F Rev | GAGCAGTTTGATCATTGATATAGAACATCATCGAGTAAAAATATGCG |
| DabA R334A For | AACGATGTGTGCACTAGCGATGCGGAAAGATTGGACGAAG |
| DabA R334A Rev | CTTCGTCCAATCTTTCCGCATCGCTAGTGCACACATCGTT |
| DabA R423A For | GTGTCGAGGATGGGTACGCTGCTGACCACAAACC |
| DabA R423A Rev | GGTTTGTGGTCAGCAGCGTACCCATCCTCGACAC |
| DabA Y143A For: | TGTCGCATATTTTTACTCGATGATGGCCTATATCAATGATCAAACTGCTCAT |
| DabA Y143A Rev | ATGAGCAGTTTGATCATTGATATAGGCCATCATCGAGTAAAAATATGCGACA |
| DabA Y143L For | GTCGCATATTTTTACTCGATGATGTTATATATCAATGATCAAACTGCTCATC |
| DabA Y143L Rev | GATGAGCAGTTTGATCATTGATATATAACATCATCGAGTAAAAATATGCGAC |
| DabA H412A For | CTTTGATTGGGTATGTATTGGCCGAAGTTTGCTGTGTCGAGG |
| DabA H412A Rev | CCTCGACACAGCAAACTTCGGCCAATACATACCCAATCAAAG |
| DabA E413A For | TGGGTATGTATTGCACGCAGTTTGCTGTGTCGAGG |
| DabA E413A Rev | CCTCGACACAGCAAACTGCGTGCAATACATACCCA |
| KabA M114T For | ACAATCGTATAAATTGTGAGAAAACGGGTTCTTTGATGGCCC |
| KabA M114T Rev | GGGCCATCAAAGAACCCGTTTTCTCACAATTTATACGATTGT |

**Table S1**. Primers for PCR

| | DabA + $Mg^{2+}$ + GSPP Complex | DabA + $Mn^{2+}$ + GSPP Complex | DabA + $Mg^{2+}$ + NGG Complex |
|---|---|---|---|
| **Accession code** | 6VKZ | 6VL0 | 6VL1 |
| **Data collection** | | | |
| Space group | $P4_32_12$ | $P4_32_12$ | $P4_32_12$ |
| Cell dimensions | | | |
| a, b, c (Å) | 124.3, 124.3, 114.5 | 124.2, 124.2, 114.0 | 123.4, 123.4, 113.1 |
| α, β, γ (°) | 90.0, 90.0, 90.0 | 90.0, 90.0, 90.0 | 90.0, 90.0, 90.0 |
| Resolution (Å) | 42.1-2.10 (2.107-2.100) | 50.0-2.20 (2.207-2.200 | 49.6-2.10 (2.107-2.100) |
| $R_{sym}$ (%) | 13.0 (121.4) | 13.2 (116.6) | 12.3 (89.8) |
| $R_{pim}$ (%) | 3.9 (35.9) | 4.6 (39.7) | 2.9 (20.0) |
| $I / \sigma I$ | 16.6 (3.2) | 15.8 (3.1) | 22.7 (6.6) |
| Completeness (%) | 94.9 (100) | 90.9 (100) | 100 (100) |
| Redundancy | 11.8 (12.3) | 9.4 (9.8) | 19.0 (20.7) |
| $CC_{1/2}$ | 0.99 (0.78) | 0.99 (0.81) | 1.00 (0.96) |
| **Refinement** | | | |
| Resolution (Å) | 42.1-2.1 | 41.0-2.2 | 42.0-2.1 |
| No. reflections | 50064 | 41668 | 51497 |
| $R_{work}$ / $R_{free}$ (%) | 16.6/18.8 | 17.8/20.3 | 17.7/19.5 |
| No. atoms | | | |
| Protein | 3581 | 3581 | 3578 |
| Water | 390 | 338 | 460 |
| *B*-factors ($Å^2$) | | | |
| Protein | 32.8 | 34.5 | 38.3 |
| Water | 42.2 | 42.2 | 46.5 |
| Ligands | 32.8 | 48.7 | 52.7 |
| R.m.s. deviations | | | |
| Bond lengths (Å) | 0.006 | 0.002 | 0.002 |
| Bond angles (°) | 0.771 | 0.415 | 0.477 |

1. Highest resolution shell is shown in parenthesis.
2. R-factor = $\Sigma(|F_{obs}|-k|F_{calc}|)/\Sigma |F_{obs}|$ and R-free is the R value for a test set of reflections consisting of a random 5% of the diffraction data not used in refinement.

**Table S2**. Refinement statistics for crystallography data.

| **Representative Terpene Cyclase Family Members Selected for Tree** | | | | **Added Sequences** |
|---|---|---|---|---|
| Q9X839 | A0A010S4I9 | A0A0A1Z8V8 | A0A0B5IKE4 | TPR00180 |
| Q9K499 | Q94G53 | J7LH11 | Q41594 | SCL16690 |
| A3KI17 | B9S9Z3 | B9RI00 | B3TPQ6 | AXN72983 |
| A0A099D720 | A0A287GX55 | A8NE23 | A0A0D3FK28 | AZO92733 |
| A0A069JMU8 | Q6Q3H2 | A0A066YQY7 | A0A061F9B8 | AZO92734 |
| B2KSJ5 | A0A0A8EZZ1 | A0A072U8B8 | A0A1S3YTU4 | ASV63464 |
| C7ASI9 | A0A060SKA0 | A0A076KZH5 | A0A1C9J6A7 | |
| A0A097CSG2 | Q82RR7 | Q84LF0 | A0A0L9V5C4 | |
| A0A097CS99 | A0A071M9D1 | A0A0J0BNS6 | A0A072UZ75 | |
| B1W019 | A0A0J8B437 | O49853 | Q84UU4 | |
| Q29VN2 | A0A098DVT4 | A0A072UXL6 | A0A067JQ70 | |
| A0A014MUN6 | B9RHP7 | C5YHH7 | A0A1S3Y8N1 | |
| A0A0D3FUB5 | A0A1J6ID15 | A0A0C1W9J4 | A0A3S3NM43 | |
| O64961 | A0A0L9TD46 | A0A0D7CID6 | P0DL13 | |
| A0A066U7L8 | A0A0C5KR55 | A0A0K8LQJ0 | A0A0B2R155 | |
| B9RXW4 | A0A0B2PNQ2 | A4FVP2 | A0A067D5M4 | |
| E2E2P0 | A0A1E5W0J5 | A0A067DG75 | Q4KSH9 | |
| B9T825 | A0A0D4DTS5 | A0A0Q3I5V4 | O48935 | |
| B0FGA9 | A0A097ZQD8 | A0A1L7U8F2 | A0A0H5CEF0 | |
| A0A0C5L205 | A0A072V8U9 | A0A0C5KH39 | A0A071M8M2 | |
| A0A067P991 | A0A1B6PFB6 | Q6PWU2 | P0CJ43 | |
| D9XD61 | A0A284QQE2 | O24475 | A0A022PQ06 | |
| B9T536 | G1JUH1 | B9SCB6 | A0A0U1XXJ7 | |
| G5CV45 | A0A059BXJ5 | B9RPM3 | P59287 | |
| Q9C6W6 | A0A059C923 | C7E5V7 | Q5SBP3 | |
| I6QPS5 | D2B747 | B2J4A4 | J7LQ09 | |
| A0A060SSS1 | E3VWJ0 | A0A0D3F4V0 | I6RAQ6 | |
| E3W208 | A0A0B2R0J5 | H8ZM70 | Q9C748 | |
| A0A077KIK5 | A0A0F4JVJ9 | A0A0I9YLV3 | A0A287XU99 | |
| A0A077RZ53 | A0A068UTJ4 | A0A072TWP0 | B6SCF5 | |
| A8NU13 | O64404 | A0A077KD00 | A0A074RGB6 | |
| A0A084T0Z4 | O81191 | A0A077KMI1 | E2E2N7 | |
| B9RXW0 | D0VMR5 | O65434 | P0DPK6 | |
| A0A0Q3E1I9 | B1B1U3 | A0A061GGG2 | A0A0J6XHN2 | |
| A0A0D3FIQ8 | A0A014L5I1 | A0A0D3FUB9 | C7PLV2 | |
| B5HDJ6 | A0A024RWJ3 | Q9FXY7 | A0A078FGP2 | |
| A0A0A9I9R9 | A0A0B2PE61 | B5A434 | A0A0A7HJU3 | |
| Q8GUE4 | A0A0B4L8B1 | E4N7E5 | Q8GUE4 | |

**Table S3**. List of NCBI accession codes for protein sequences used to construct the maximum likelihood phylogenetic tree. 150 were selected as representatives from the InterPro and 6 additional sequences were added as described on pg S5.

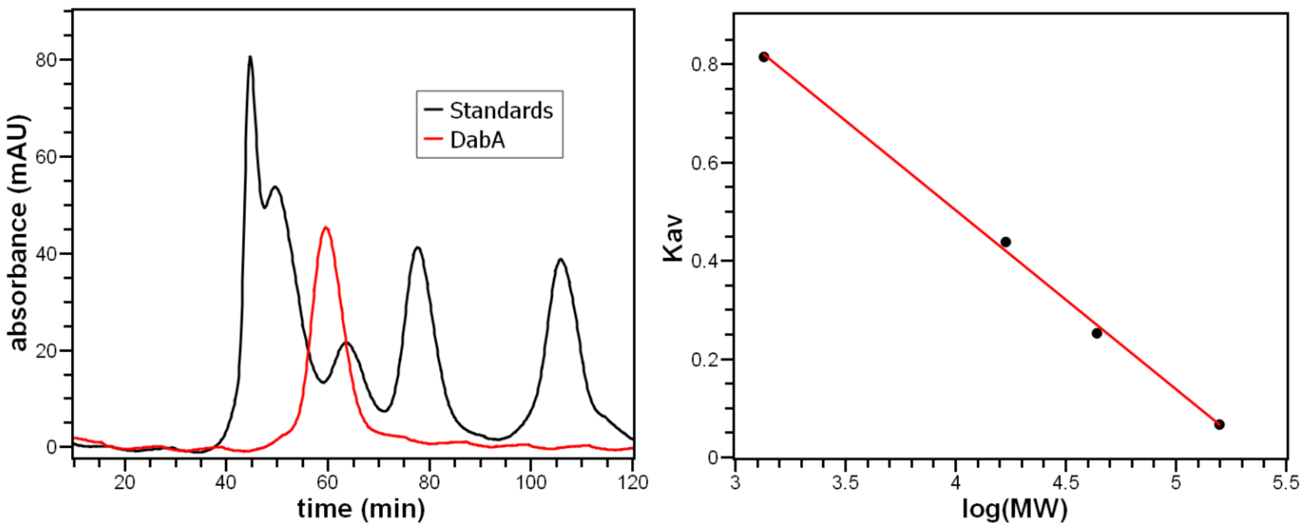


**Figure S1**: Gel filtration was used to estimate oligomerization state of DabA in solution. A standard curve was generated using BioRad Gel Filtration Standard (#1511901) and the experiment was completed on a HiLoad 16/60 Superdex 75 prep grade column. DabA eluted at 60 mins and was estimated to be 68.6 kDa. The molecular weight of the $His_6$-tag cut DabA Glu46 to C-term construct is 51.5 kDa. The calculated solution state is 1.3 monomers in size, and, therefore, DabA likely exists a monomer.

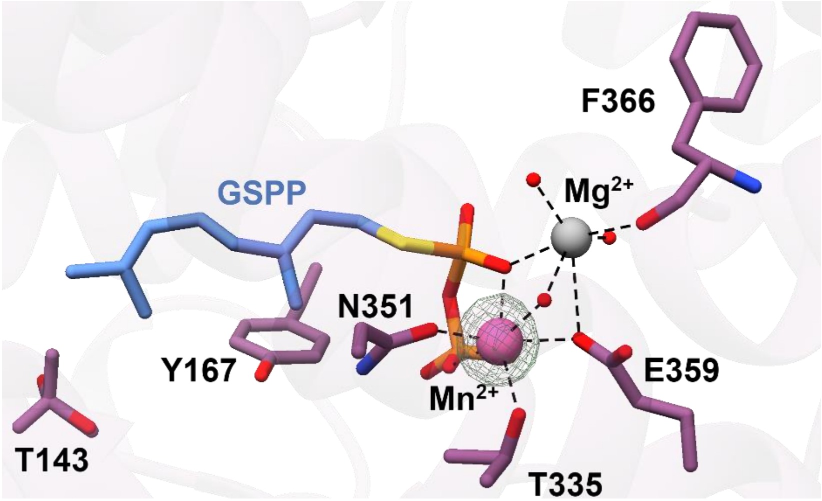


**Figure S2**: Active site of DabA in complex with GSPP. Crystals were soaked in a solution containing $MnCl_2$ in order to replace magnesium ions with the more electron dense manganese. $F_o$-$F_c$ maps were generated by removing the manganese and completing refinement without the ion. Mesh was contoured to 10σ (green). Based on the observed electron density, manganese could successfully substitute for one of the two magnesiums.

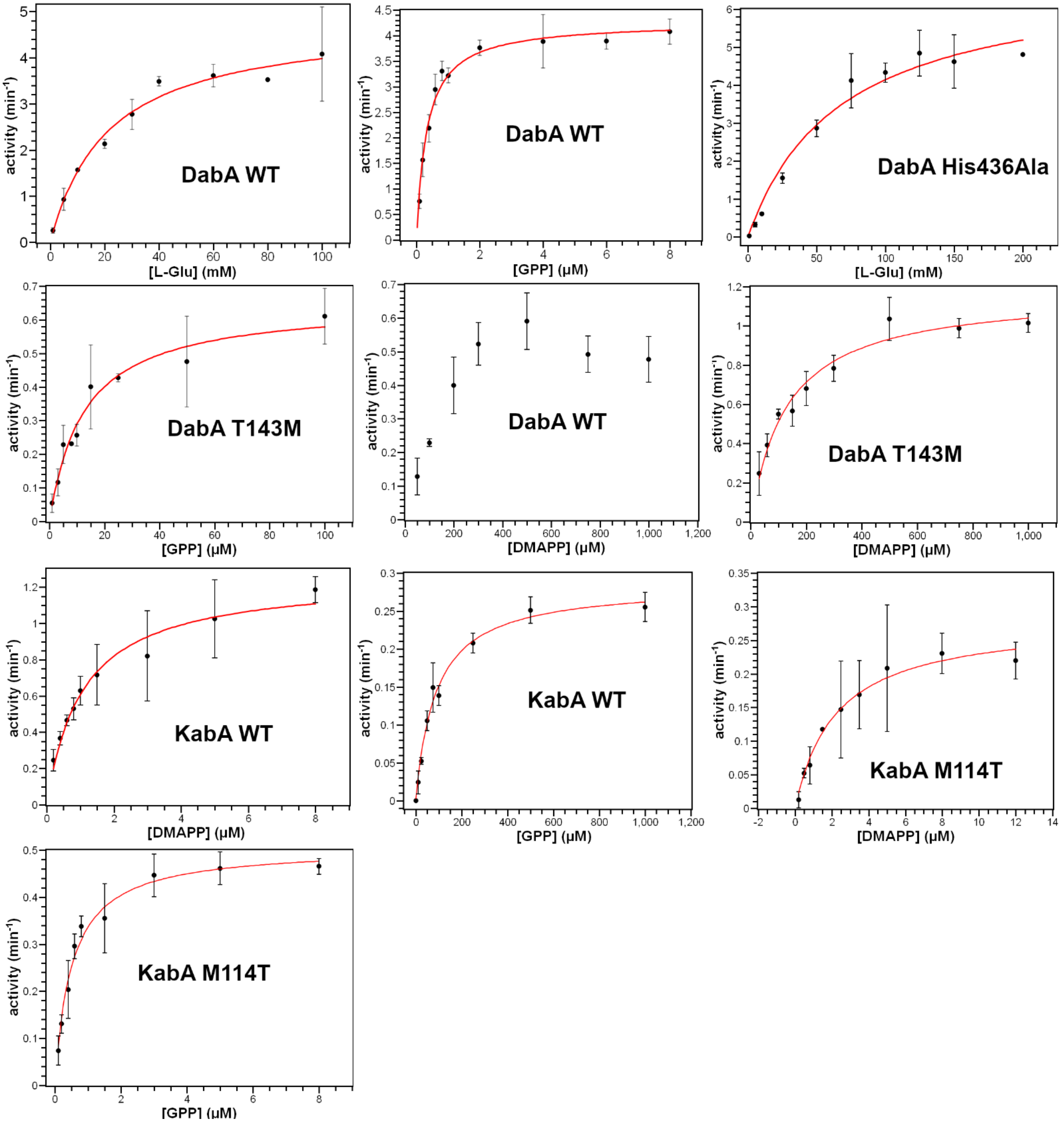


**Figure S3**. Kinetic analysis of DabA, KabA and variants. Data was fit to the Michaelis-Menten equation. Each point was completed in triplicate. Values for kinetic parameters are found in Table 1. DabA did not display Michaelis-Menten kinetics when DMAPP was used as a substrate. This data was also fit to the substrate inhibition Michaelis-Menten equation, but the resulting constants were not physiologically relevant.

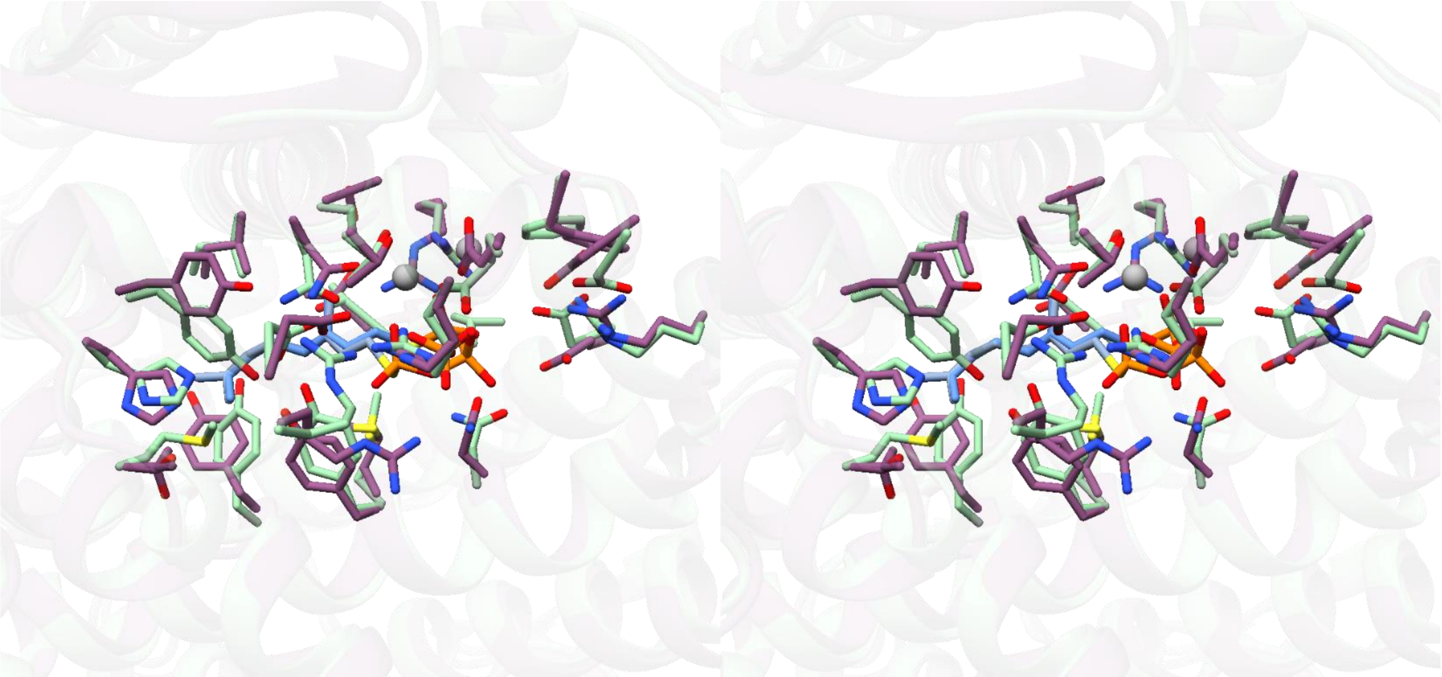

**Figure S4**: Stereoview of the DabA active site (purple) aligned with the KabA I-TASSER model (green). GSPP is shown in blue.

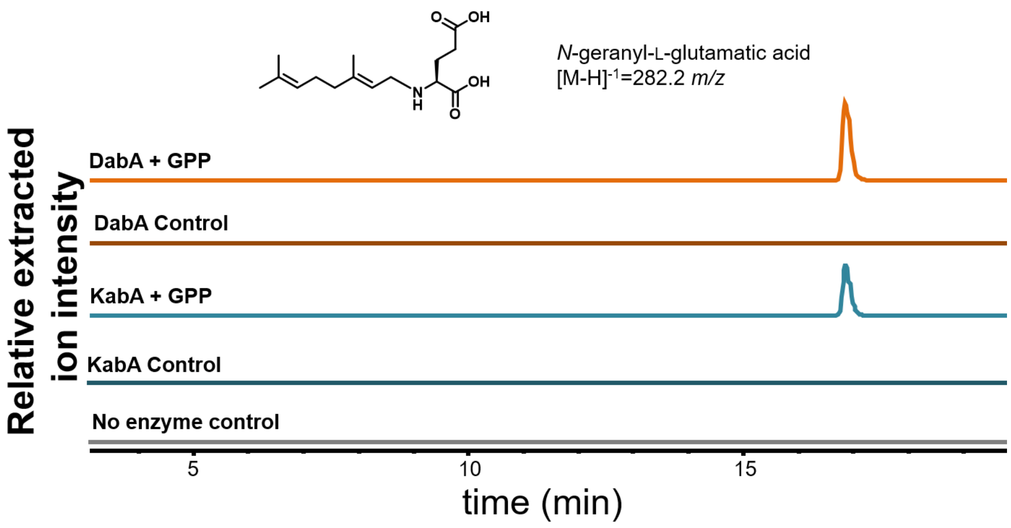

N-geranyl-L-glutamatic acid
[M-H]-1=282.2 m/z
Relative extracted ion intensity
DabA + GPP
DabA Control
KabA + GPP
KabA Control
No enzyme control
5
10
15
time (min)


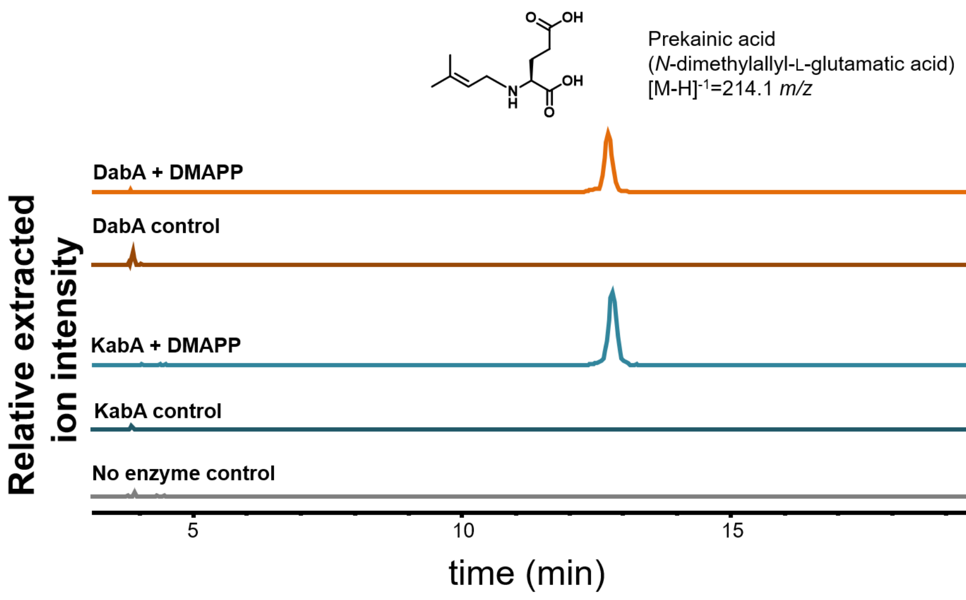

Prekainic acid
(N-dimethylallyl-L-glutamatic acid)
[M-H]-1=214.1 m/z
Relative extracted ion intensity
DabA + DMAPP
DabA control
KabA + DMAPP
KabA control
No enzyme control
5
10
15
time (min)


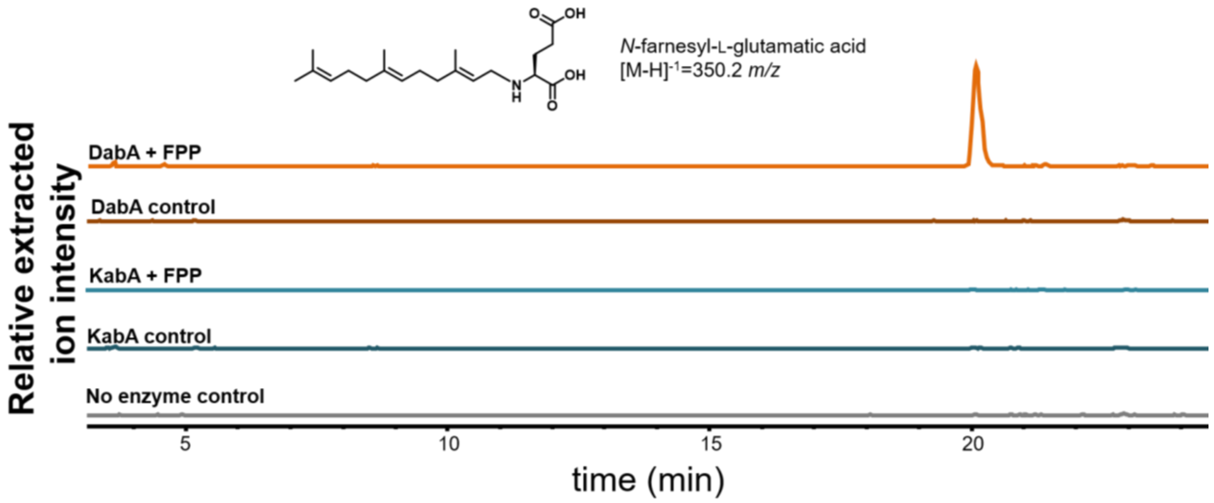

N-farnesyl-L-glutamatic acid
[M-H]-1=350.2 m/z
Relative extracted ion intensity
DabA + FPP
DabA control
KabA + FPP
KabA control
No enzyme control
5
10
15
20
time (min)

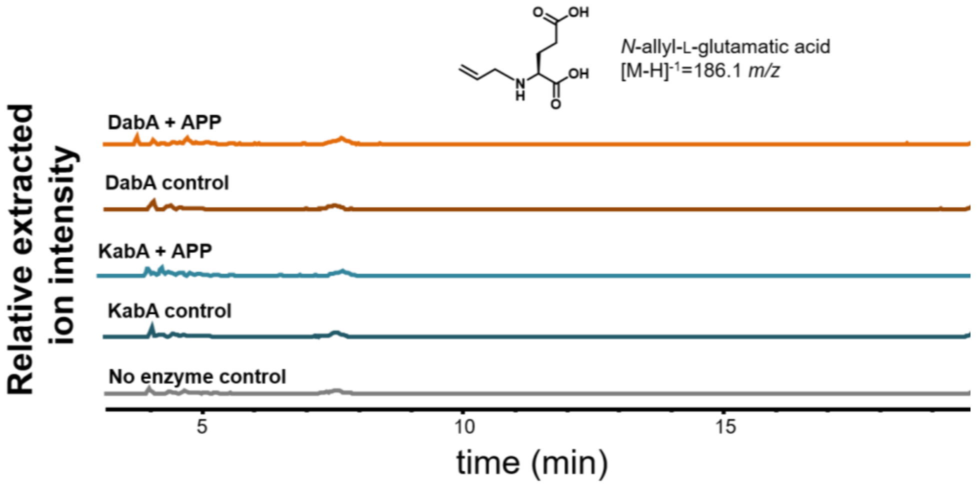

*N*-allyl-L-glutamatic acid
[M-H]-1=186.1 *m/z*
Relative extracted ion intensity
DabA + APP
DabA control
KabA + APP
KabA control
No enzyme control
5
10
15
time (min)


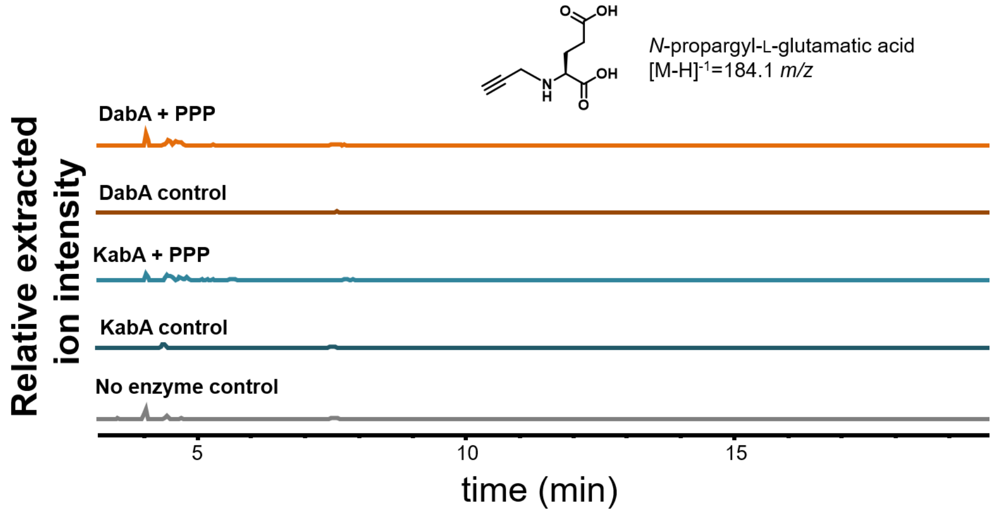

*N*-propargyl-L-glutamatic acid
[M-H]-1=184.1 *m/z*
Relative extracted ion intensity
DabA + PPP
DabA control
KabA + PPP
KabA control
No enzyme control
5
10
15
time (min)


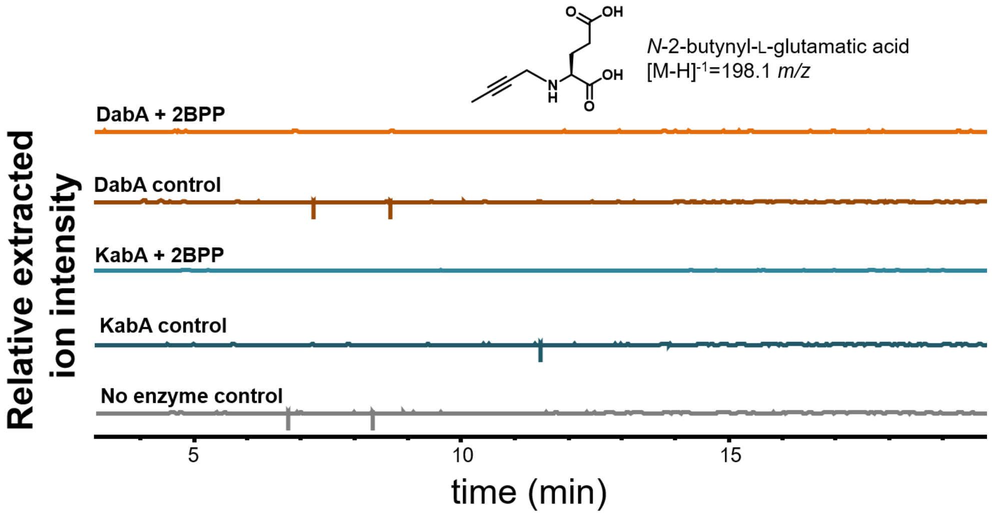

*N*-2-butynyl-L-glutamatic acid
[M-H]-1=198.1 *m/z*
Relative extracted ion intensity
DabA + 2BPP
DabA control
KabA + 2BPP
KabA control
No enzyme control
5
10
15
time (min)

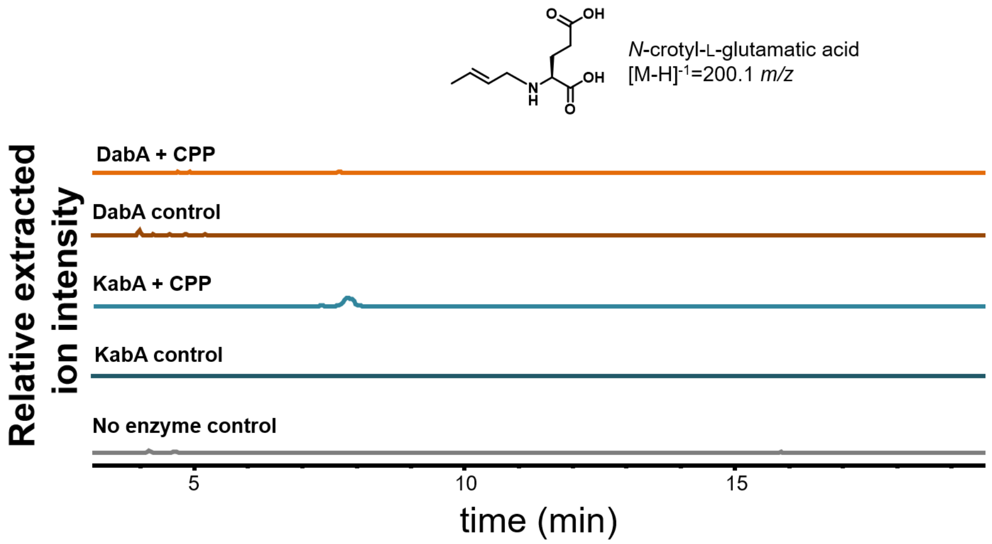


**Figure S5**: Both KabA and DabA were tested with a variety of prenyl diphosphates to evaluate their substrate specificity. Each trace is the EIC (extracted ion chromatogram) ± 0.5 *m/z* for the expected mass of the product indicated. The "DabA and KabA controls" lacked the prenyl diphosphate while the "No enzyme control" lacked enzyme. Substrate abbreviations are as follows: GPP (geranyl diphosphate), DMAPP (dimethylallyl diphosphate), FPP (farnesyl diphosphate), PPP (propargyl diphosphate), 2BPP (2-butynyl diphosphate), APP (allyl diphosphate), and CPP (crotyl diphosphate).

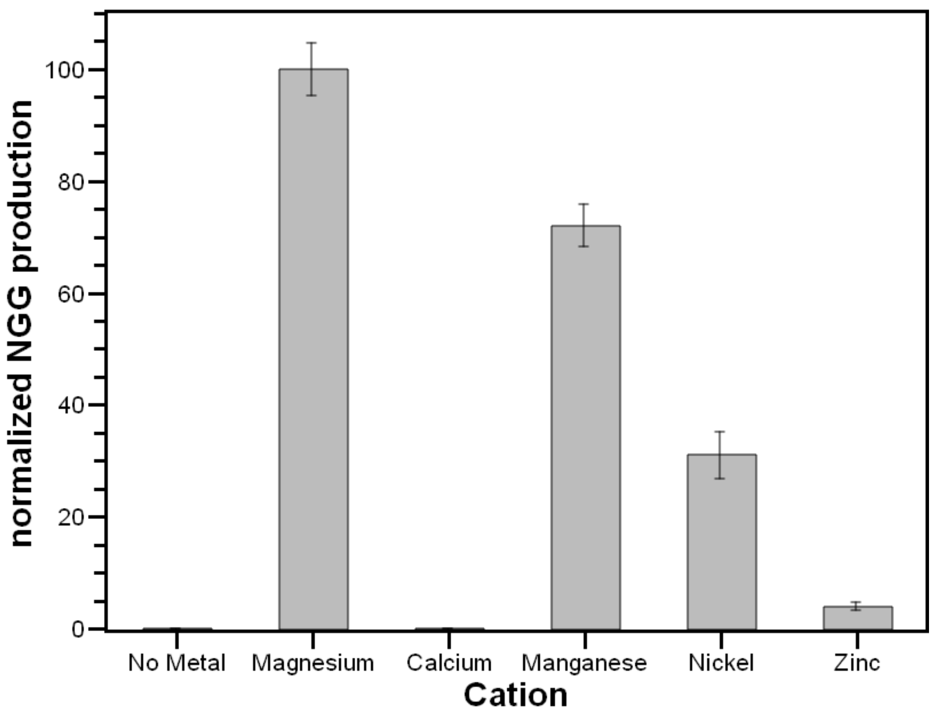


**Figure S6:** DabA activity assays were completed with either $MgCl_2$, $CaCl_2$, $MnCl_2$, $NiCl_2$, $ZnCl_2$, or no added metal to evaluate the importance of the identity of the cation for catalysis. While several metals can be used for catalysis, magnesium appears to be the favored divalent cation.
